# Insular error network enables self-correcting intracranial brain-computer interface

**DOI:** 10.1101/2025.11.17.688824

**Authors:** Paul Weger, Maarten C. Ottenhoff, Maxime Verwoert, Sophia Gimple, Lauren Ostrowski, Albert Colon, Louis Wagner, Johannes P. van Dijk, Yasin Temel, Pieter Kubben, Christian Herff

## Abstract

Error recognition is fundamental to adaptive behavior, enabling rapid compensatory action when outcomes deviate from expectations. Central to this function are neural circuits for performance monitoring, encoding cognitive signals that could support more reliable neural interfaces. Here, we recorded intracranial electroencephalography (iEEG) in epilepsy patients to enable a motor brain-computer interface (BCI) while sampling error-related activity across a distributed network. Our work reveals high-frequency population dynamics emerging in the anterior insula and propagating to the prefrontal cortex as the interface fails to follow the user’s intention. We identify spatially organized insular responses to error processing and movement feedback, highlighting it as a heterogeneous hub linking action and outcome. Real-time integration of error responses enables a self-correcting neural interface that enhances usability by reducing the need for manual user intervention. Together, our work demonstrates a human intracranial BCI harnessing insular brain activity, integrating cognitive processes directly into device control.

## Introduction

Recognizing one’s error is an essential function that allows behavior to remain aligned with goals. When the brain detects a mismatch between an intended action and its outcome, it generates a characteristic neural response known as an Error-related potential (ErrP). These signals were first identified in the early 1990s using electroencephalography (EEG) and are typically characterized by a fronto-central negativity followed by delayed positivity^1,2^.

Investigating the neural mechanisms underlying ErrPs offers insight into how the brain monitors performance and adapts behavior. Studies have therefore explored the neural correlates of error processing using a wide range of methods, spanning non-invasive techniques such as EEG and functional magnetic resonance imaging (fMRI), to invasive intracranial recordings of spiking activity and Local Field Potentials (LFPs). Converging evidence points to widespread cortical–subcortical circuits, with the salience network and the dopaminergic system playing central roles in detecting mismatches and driving behavioral adaptation^3–6^.

Within these circuits, several anatomical regions have been identified as key hubs for error detection, most notably the dorsal anterior cingulate cortex (dACC)^7–10^ and anterior insula (aINS)^11–16^. These regions are thought to dynamically regulate behavior by signaling the need for adjustment and driving widespread cortical responses. Following the detection of an error, other brain areas contribute to coordinating and initiating corrective actions. Research suggests involvement of the prefrontal cortex (PFC) in supporting higher-order cognitive adjustments^10,17,18^, while central and parietal regions contribute to integrating sensory feedback and updating motor plans^8,19,20^.

For individuals with severe motor impairments, such as those with spinal cord injury, amyotrophic lateral sclerosis (ALS), or locked-in syndrome, brain-computer interfaces (BCIs) offer a way of regaining control and interacting with the environment^21,22^. However, unreliable decoding of neural activity can lead to a loss of confidence and trust in the system, limiting its long-term effectiveness. Integrating error-related potentials into BCI control could enable automatic detection and correction of mistakes, thereby enhancing reliability and user satisfaction^23,24^.

Error-related potentials during BCI usage typically arise when a decoder misclassifies a neural signal and triggers an unintended action^24,25^. Their presence in non-invasive BCIs is well established, with both offline and online decoding approaches reliably distinguishing correct from erroneous trials^23,26^. Building on this work, invasive studies in non-human primates (NHPs) have demonstrated robust offline error encoding in primary motor cortex (M1) and premotor cortex (PM) during brain-machine control tasks^15,27^. More recently, these findings have been extended to humans, with intracranial recordings revealing similar error-related signals in the hand-knob area and speech motor cortex^28,29^.

Substantial research has characterized error-related potentials offline during BCI use, providing a basis for tracking these signals in real time. Online error correction builds on this approach by identifying ErrPs and using them to automatically adjust or override BCI outputs. Its feasibility has been demonstrated in EEG-based BCIs, where integrating error correction reliably improved control accuracy^30–32^. Invasive studies in NHPs expanded on this work by implementing both trial-end corrections and continuous error tracking during two-dimensional cursor movements and multi-finger hand control^33,34^.

While previous studies establish the feasibility of decoding error signals for adaptive control, their application in human intracranial BCIs remains largely unexplored. To address this gap, we recorded intracranial electroen-cephalography from epilepsy patients as they controlled a motor BCI, in which grasping movements were used to steer a car-racing game. The broad cortical–subcortical coverage of iEEG offered a rare window into deep and distributed nodes of the error-processing network, enabling precise spatiotemporal tracking of neural activity for automatic error detection and correction (Fig. 1a).

**Figure 1.**
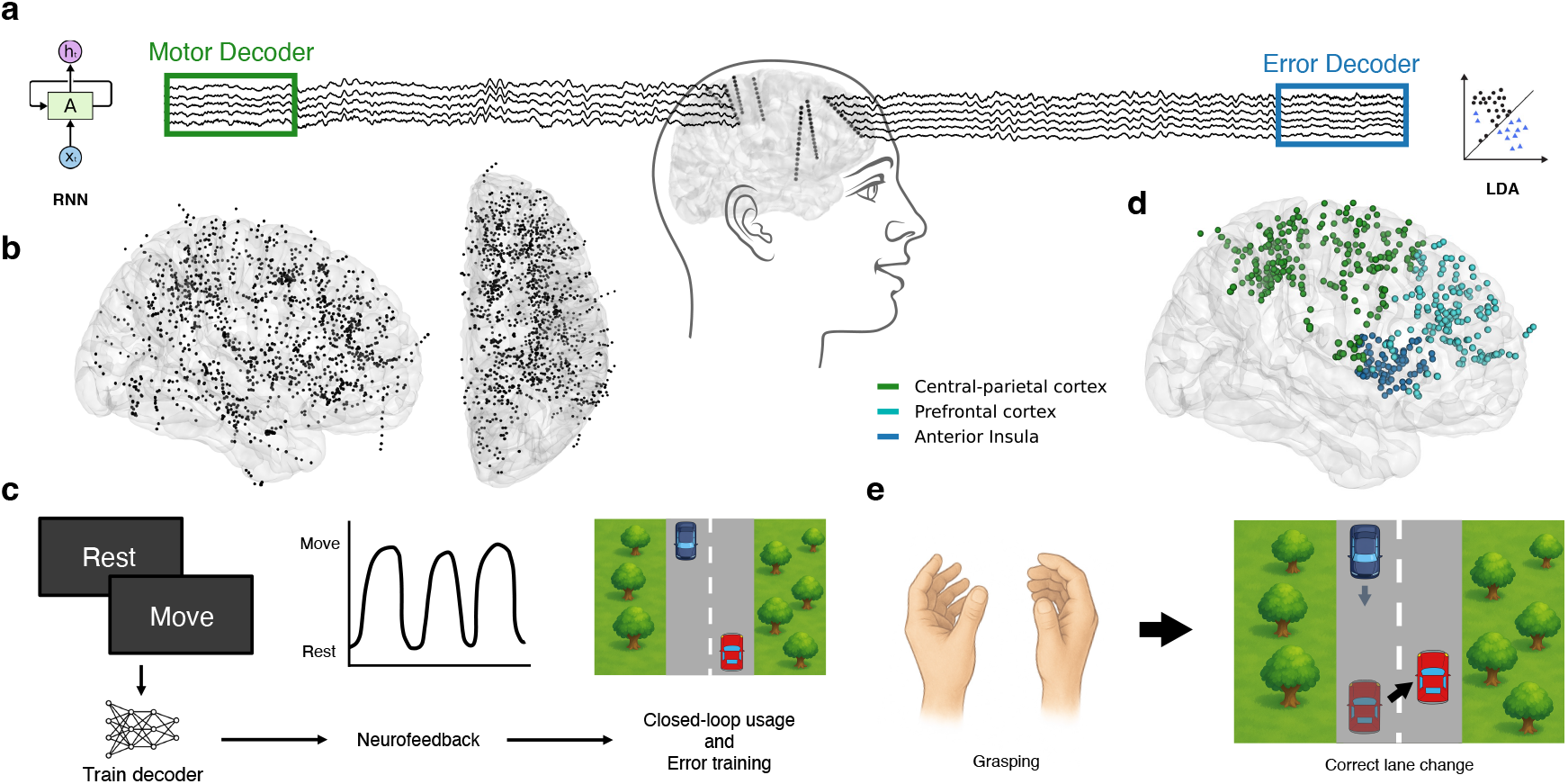
Schematic of the task paradigm and electrode coverage. (**a**) Example participant with electrodes in motor-relevant regions and prefrontal areas central to error-related activity. Two decoders operated in parallel to enable error correction during motor BCI control. (**b**) Recording sites from all 10 participants, warped to the Montreal Neurological Institute (MNI) average brain for visualization. (**c**) Overview of the three-stage task paradigm: a calibration task to train the movement decoder, a neurofeedback session to foster patient adaptation, and a racing game to test performance and track error-related activity. (**d**) Regions of interest included in the analysis of error-related potentials during closed-loop BCI use. (**e**) Game control of the racing game, where grasping triggers a lane change of the red player car to avoid oncoming obstacles.

## Results

### Participants and task paradigm

We analyzed iEEG recordings from 10 participants (6 female, age range 26-56 years) undergoing clinical monitoring for drug-resistant epilepsy. Electrode implantation was solely determined by clinical needs, providing access to 1,317 channels across cortical and subcortical structures (Fig. 1b). Analysis of error-related activity focused on three regions of interest: the anterior insula (73 electrodes), prefrontal cortex (160 electrodes), and a combined set of central and parietal implantation sites (242 electrodes, Fig. 1c).

Participants started the three-stage task paradigm by performing a movement calibration task for three minutes (Fig. 1d). They alternated between resting and grasping movements, cued by a visual countdown bar, to generate training data for a real-time movement decoder. Next, patients adapted to the model’s output through neurofeedback, during which they observed decoder predictions for two minutes while performing self-paced grasping. This familiarization period helped patients adjust to the system and better anticipate its outputs. Finally, they played a computerized racing game, in which grasping movements triggered a lane change to avoid oncoming cars (Fig. 1e). The first round of the game was used to track neural responses to errors, providing data to train a second decoder running in parallel to the movement decoder (Fig. 1a).

### Brainwide spectrotemporal signatures of movement

Calibrating the motor BCI required only three minutes of supervised data, during which participants followed task instructions shown on a screen (Fig. 2a). Neural activity was captured in two key frequency ranges of movement: high-frequency activity (HFA, 60–200 Hz), and low-frequency activity (LFA, 8–30 Hz), corresponding to alpha and beta rhythms. HFA is commonly associated with local neuronal firing, whereas LFA captures broader oscillatory dynamics associated with motor activity^35^. As expected, we observe a sharp rise in HFA and a simultaneous reduction in LFA at movement onset. Power in both frequency bands showed a transient peak following the grasping instruction, with HFA reaching its maximum at 520 ms and LFA reaching its minimum at 540 ms (Fig. 2c). After the peak, neural power stabilized during sustained movement, reaching a plateau at 1.15 s for high frequencies and 1.04 s for low frequencies.

**Figure 2.**
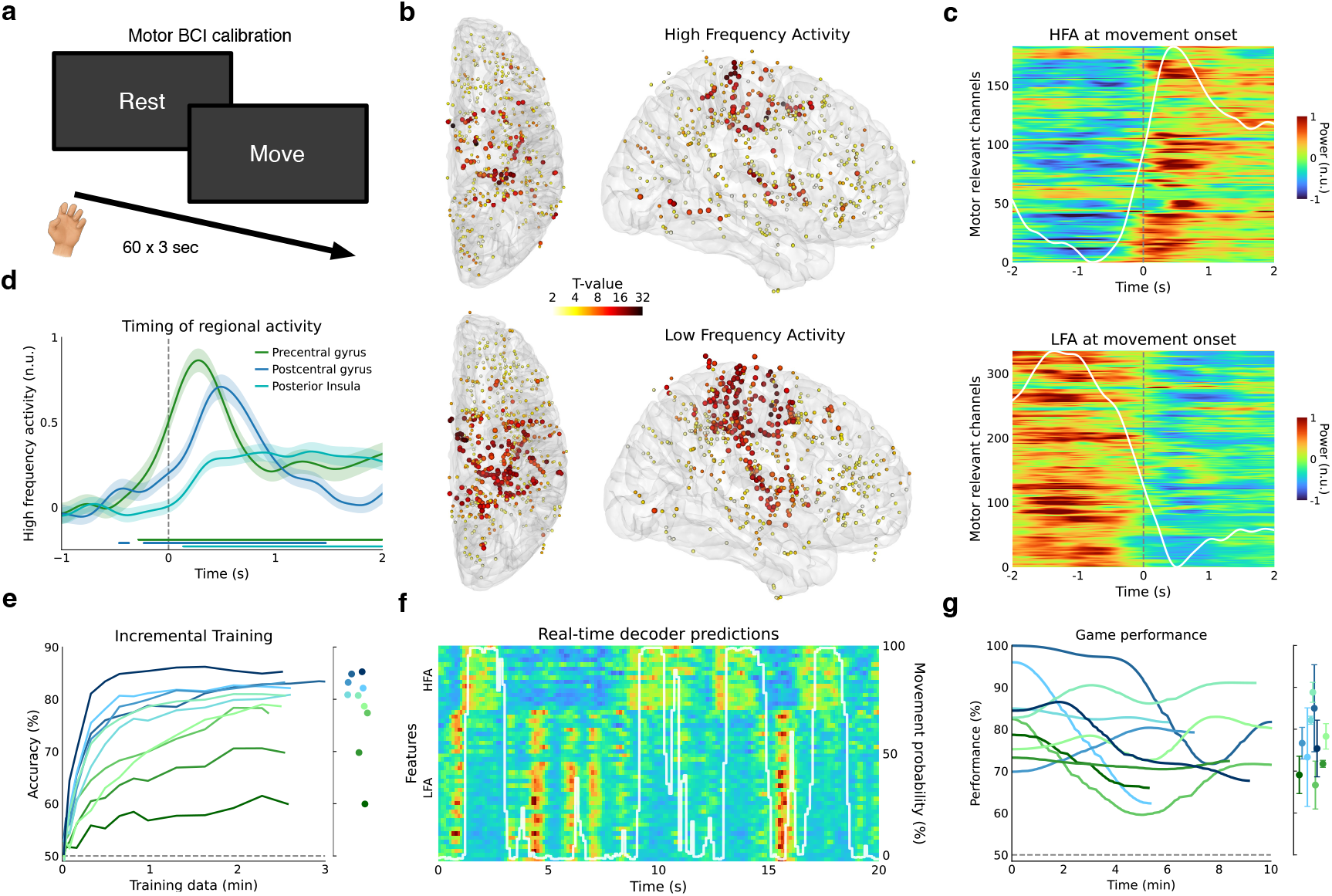
Spatial, spectral, and temporal features underlying movement and decoder training a, Calibration task with visual cues and task structure. **b**, Spatial distribution of significantly movement-related electrodes, shown separately for high-frequency activity (top panels) and low-frequency activity (bottom panels). All electrodes have been mapped onto one hemisphere. **c**, Population-level neural activity from highly movement-related channels in high-frequency and low-frequency bands (*t >* 4, *p <* 0.05). The grey dashed line marks promoted movement onset (0); white trace shows the mean across channels. **d**, Regional timing of HFA around prompted movement onset for three motor-active areas. Bold lines represent averages across electrodes, shaded areas indicate standard error of the mean (s.e.m.). Colored bars mark time points significantly different from baseline (HFA during resting, –2 to –0.5 s before movement onset). **e**, Incremental decoder training as a function of calibration data length, from 1 second up to the full 3 minutes. The right panel shows the final accuracy used for BCI control. Each participant is shown in a different color; light grey dashed line indicates chance level. **f**, Example of real-time decoder output (white) and contributing spectral features (HFA/LFA, colored) during gameplay in an exemplary participant. **g**, Game performance across participants over time, colored as in **e**; average performance shown on the right panel.

The spatial distribution of slow and fast neural dynamics revealed high-frequency changes clustering around central regions, with responsive electrodes in the pre- and postcentral gyri and a few in the supplementary motor area (Fig. 2b, upper panels). In contrast, LFA decreases exhibited a broader spatial distribution, spreading outward from the central sulcus into parietal regions and secondary motor areas, including premotor and supplementary motor cortices (Fig. 2b, lower panels). Notably, power in both frequency bands reliably activated electrodes in the insula, with LFA engaging the region to a wider extent.

To characterize the sequential engagement of motor-related regions during movement initiation, we analyzed the timing of HFA in the precentral gyrus (including the primary motor cortex), postcentral gyrus (including the primary somatosensory cortex), and posterior insula. Activity at movement initiation peaked earliest and strongest in the precentral gyrus (275 ms after prompted movement onset, HFA of 0.86 n.u.), followed by delayed peaks with smaller amplitude in the postcentral gyrus (480 ms, 0.71 n.u.) and posterior insula (first peaking at 415 ms with a power of 0.29 n.u., Fig. 2d). A continuous period of significantly increased HFA relative to baseline power (*p <* 0.05) was first observed in the precentral gyrus (280 ms before cued instruction until 2 s afterwards), and subsequently in the postcentral gyrus (−230 ms to 1.42 s) and posterior insula (140 ms to 2 s). Activity in central areas declined more rapidly after the initial peak, reaching 36% and 11% of peak amplitude in the precentral and postcentral gyri two seconds after prompted movement initiation. In contrast, posterior insula activity remained higher during ongoing movement at 93% of its peak amplitude after two seconds.

Subsequently, we trained a recurrent neural network (RNN) decoder using spectral features from both low- and high-frequency bands, allowing the model to integrate information across regions and timescales. All participants achieved decoding accuracies above chance level (50%), ranging from 59% in the lowest-performing individual to 86% in the highest-performing individual (Fig. 2e, right panel). Importantly, 8 out of 10 participants reached accuracies exceeding 75%. Incremental training with increasing amounts of calibration data revealed rapid improvements within the first minute, accounting on average for 83% of the improvement from chance level to final accuracy (Fig. 2e). This trend was followed by plateauing decoding performance after using two minutes of training data, reaching on average 98% of the top accuracy. Once trained, the decoder was deployed in real-time to control the closed-loop BCI task. The model’s output closely reflected the underlying spectral dynamics during gameplay, with rises in HFA reliably preceding increases in movement probability in an exemplary patient (Fig. 2f). LFA, in contrast, exhibited movement-related power decreases but showed greater variability across channels and time, making it a less reliable feature.

Game performance varied across participants, with average game success exceeding 65% in all individuals (Fig. 2g, right panel). Notably, even participants with calibration accuracies as low as 60% were able to control the car above chance level (50%) throughout the entire session (Fig. 2g, left panel).

Together, we demonstrate a real-time brain-computer interface that leverages low- and high-frequency dynamics to enable reliable game control. We take advantage of the brain-wide coverage of iEEG to sample distributed motor areas, showing sequential activation at movement onset and highlighting the posterior insula as a distinct and persistent feature during ongoing movement.

### Time-delayed error processing in insular-prefrontal error circuits

Next, we focused on neural responses during closed-loop BCI sessions to better understand the neural basis of error processing. We identified two possible game scenarios during real-time car control: correct lane changes, where participants successfully avoided the obstacle by moving into the free lane, and erroneous lane changes, where the car instead moved into the obstacle lane (Fig. 3a). As participants played the game with the goal of avoiding collisions, correct lane changes reflected intended decoder control and were preceded by grasp-related neural activity. In contrast, erroneous lane changes mostly arose from decoder misclassifications or unintended activations and occurred without intentional grasping. These unexpected lane changes violated participants’ expectations, creating a mismatch between the intended and actual outcomes, and were thus perceived as errors.

**Figure 3.**
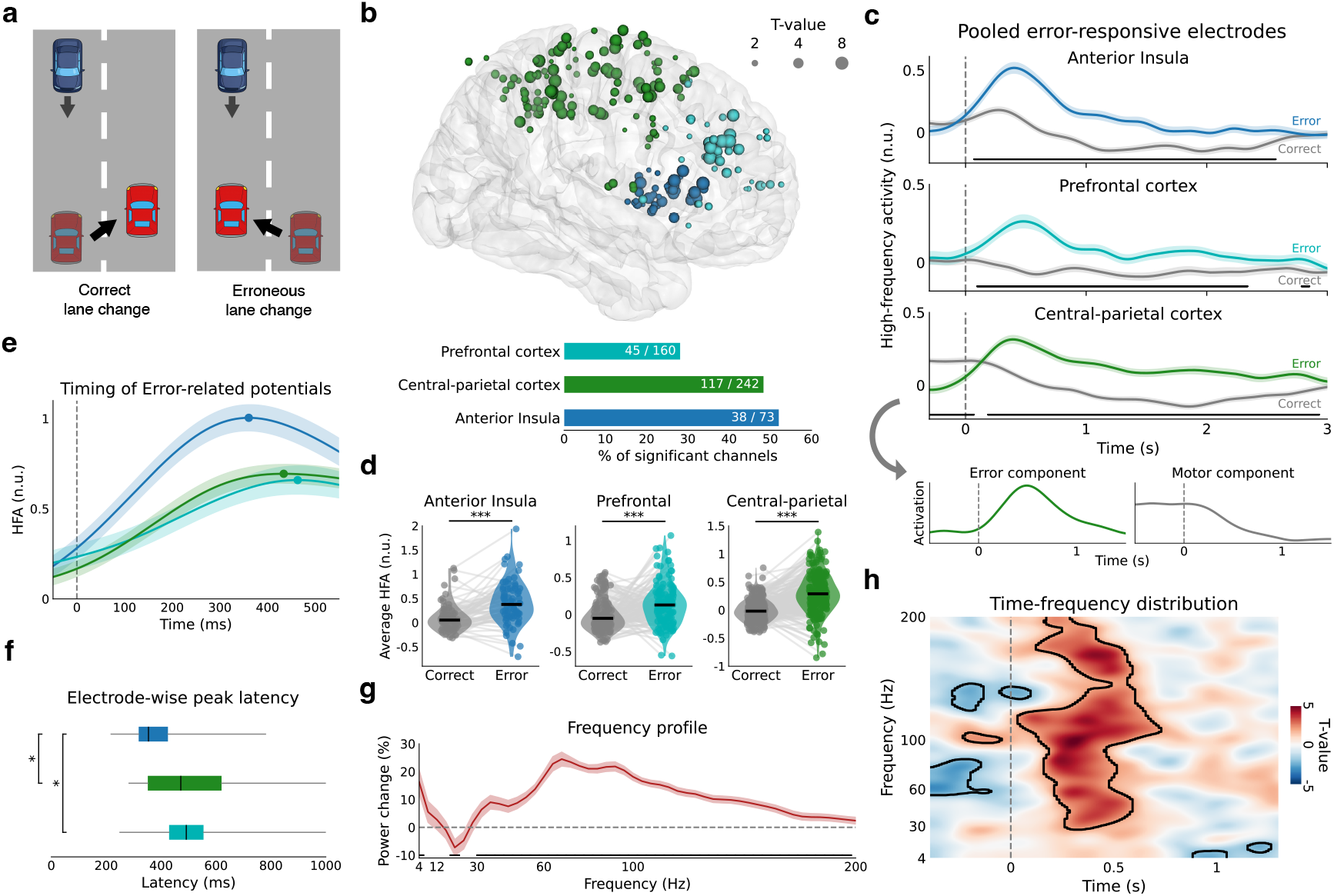
Error-related activity spreading from the anterior insula. **a**, Game scenarios during closed-loop BCI sessions. Lane changes were categorized as correct when participants avoided the obstacle by switching into the free lane, and as erroneous when they moved into the obstacle lane. **b**, Spatial distribution of significantly error-responsive electrodes across the three target regions. T-values, calculated from HFA within the first second after the lane change, are logarithmically mapped to electrode size. Bar plots in the lower panel show the percentage of significant electrodes per region, with the absolute counts (significant/total) indicated in white. **c**, Temporal trajectories of error responsive electrodes (*p <* 0.05) across three target regions. Colored lines indicate average HFA during erroneous trials, shaded areas denote s.e.m., and grey traces show correct trials. Stimulus onset is marked by a dashed line at *t* = 0, black bars indicate significance (*p <* 0.05). For the central–parietal cortex, the first two ICA components are shown, illustrating the presence of an error and movement component in this region. **d**, Power increases within the first second after stimulus presentation across all electrodes in a region. Black line marks the median of each distribution. All distributions are significantly different (*p <* 0.001, Linear mixed-effects model) between the two conditions. **e**, Trial-averaged HFA (mean ± s.e.m.) of highly responsive electrodes (*t >* 4, *p <* 0.05), with peak power marked by a dot. **f**, Peak latency of highly responsive electrodes, shown as boxplots indicating median and interquartile range. Significant pairwise differences: aINS vs. PFC (*p <* 0.05), aINS vs. central–parietal cortex (*p <* 0.05); PFC vs. central–parietal cortex was not significant (*p >* 0.05, n.s.). **g**, Frequency profile of highly responsive electrodes showing power change across frequencies, with a strong gamma/high-gamma increase. **h**, Time–frequency distribution of a representative anterior insula electrode, shown as a *t*-valued spectrogram. Significant clusters are outlined in black (*p <* 0.05).

We found error-responsive channels in the prefrontal cortex, anterior insula and central-parietal cortex (Fig. 3b), whose contribution we quantified by computing t-values of HFA over the first second following errors (unpaired two-sample t-test, *p <* 0.05). In the PFC, most responsive electrodes were clustered in the dorsolateral region (dlPFC; Brodmann areas 9 and 46), with additional electrodes more sparsely distributed across other frontal areas. Activity in the anterior insula was more concentrated in the dorsal anterior portion, where electrodes tended to show stronger responses. In contrast, channels in the central–parietal cortex were widely distributed and lacked a clear focal pattern. Overall, significant activity was observed in 28% of channels in the PFC (45/160), 48% in the central–parietal cortex (117/242), and 52% in the anterior insula (38/73).

Temporal dynamics at the population level of responsive electrodes revealed a rapid rise in HFA following erroneous lane changes in the anterior insula (Fig. 3c, top panel), peaking at 0.52 n.u. after 400 ms before gradually returning to baseline (0 n.u.) by 1.97s. Correct trials showed a different pattern, with only a modest initial peak, followed by a sustained reduction in HFA that reached a minimum of −0.15 n.u. at 1.52 s, before returning to baseline after 2.68 s. The two powers remained significantly different from each other for over 2 s, starting from 70 ms to 2.5 s (unpaired two-sample t-test, *p <* 0.05).

Prefrontal activity followed a similar pattern, with a slower initial rise after erroneous lane changes, peaking at 480 ms (0.64 n.u.) and returning to baseline by 2.85 s (Fig. 3c, middle panel). During correct trials, HFA dropped below baseline, reaching a minimum power of −0.1 n.u. at 1.45 s and remaining reduced until 1.85 s. As a result, erroneous and correct trials remained significantly separable for a continuous period of 2.2 s, starting at 95 ms and ending at 2.34 s.

In central and parietal cortices, erroneous trials showed comparable dynamics to the PFC, peaking at 400 ms (at a power of 0.31 n.u.) and returning to baseline by 3 s (Fig. 3c lower panel). Correct trials, in contrast, displayed a distinctively different pattern: activity was elevated before the lane change (0.2 n.u.), dropped below baseline 720 ms after the event, and remained suppressed thereafter. The two conditions thus remained significantly different over two intervals: from 350 ms before to 70 ms after the lane change, and from 185 ms to 2.9 s after the stimulus.

Evidently, high-frequency activity in the central–parietal cortex was elevated prior to a lane switch in correct trials, whereas it remained near baseline in the prefrontal cortex and anterior insula. This pattern seems to align with the task layout, in which correct lane switches were preceded by movement to avoid the oncoming obstacle, while erroneous switches occurred mostly without prior activity. Our results might therefore highlight that HFA in the central–parietal cortex codes both pre-switch movement and post-switch error-related activity (lowest panel in Fig. 3c).

Across all electrodes within each region, high-frequency activity significantly increased during erroneous trials compared to correct trials (Linear mixed effects model, *p* < 0.001, Fig. 3d). This increase was most pronounced in the anterior insula, where average HFA rose from 0.05 n.u. during correct trials to 0.38 n.u. during erroneous trials (z-scored power values). In the prefrontal cortex, HFA increased from –0.05 n.u. to 0.13 n.u., while power in the central-parietal cortex rose from −0.02 n.u. to 0.29 n.u.

To analyzing the precise timing of peak responses, we focused on channels with strong and reliable error-related responses (*t* > 4, *p* < 0.05), excluding those with marginal effects. We find HFA peaking at 361 ms (1 n.u.) in the anterior insula, and later at 434 ms (0.69 n.u.) in the central-parietal cortex and 463 ms (0.66 n.u.) in the prefrontal cortex (Fig. 3e). Electrode-wise peak latencies demonstrated a comparable temporal pattern, with median latencies of 354 ms in the anterior insula, 472 ms in central-parietal cortices, and 492 ms in the prefrontal cortex (Fig. 3f). Latencies in both the central–parietal (*p* < 0.05) and prefrontal (*p* < 0.05) regions were significantly delayed relative to the anterior insula, whereas no significant difference was found between the central–parietal and prefrontal cortices. Together, these temporal differences indicate that the anterior insula consistently activates earliest within the error-processing network.

Spectral power changes in highly responsive electrodes during erroneous trials revealed a distinct oscillatory signature, with elevated activity in the gamma and high-gamma range, peaking at 68 Hz (Fig. 3g). Smaller peaks were observed in the low-frequency range at individual frequency bins, with higher power at 4 Hz and a reduction at 20 Hz. Statistically different frequency bins (two-tailed one-sample t-test vs. session-wide baseline, *p* < 0.05) were observed between 32 and 196 Hz as well as 4 and 20 Hz. In an exemplary anterior insula electrode (Fig. 3h), significantly enhanced gamma/high-gamma activity (30–200 Hz) emerged 50 ms after stimulus presentation and persisted until 750 ms post stimuli.

Together, these results define the spatial, temporal, and spectral characteristics of error-related potentials during real-time BCI use. High-gamma activity in the anterior insula consistently occurred before responses in the dorsolateral PFC and central-parietal cortex, identifying it as an early node in the error-processing network.

### Distributed coding of movement and error in the human insula

To better understand the insula’s role in movement control and error monitoring, we examined its functional contributions in greater detail. We analyzed activity from 129 insular recording sites, with 56 electrodes in the posterior part and 73 in the anterior part (Fig. 4a). We examined movement-related activity using the bilateral grasping task used for decoder calibration, revealing that responsive electrodes were found primarily in the posterior and central portions of the insula, clustering near the central insular sulcus with some spread across adjacent regions (Fig. 4b). When projected along the posterior–anterior axis, the distribution of significant contacts (shown in blue) displays a clear peak in the long gyri of the posterior insula, with a second, smaller peak in the central portion of the anterior insula.

**Figure 4.**
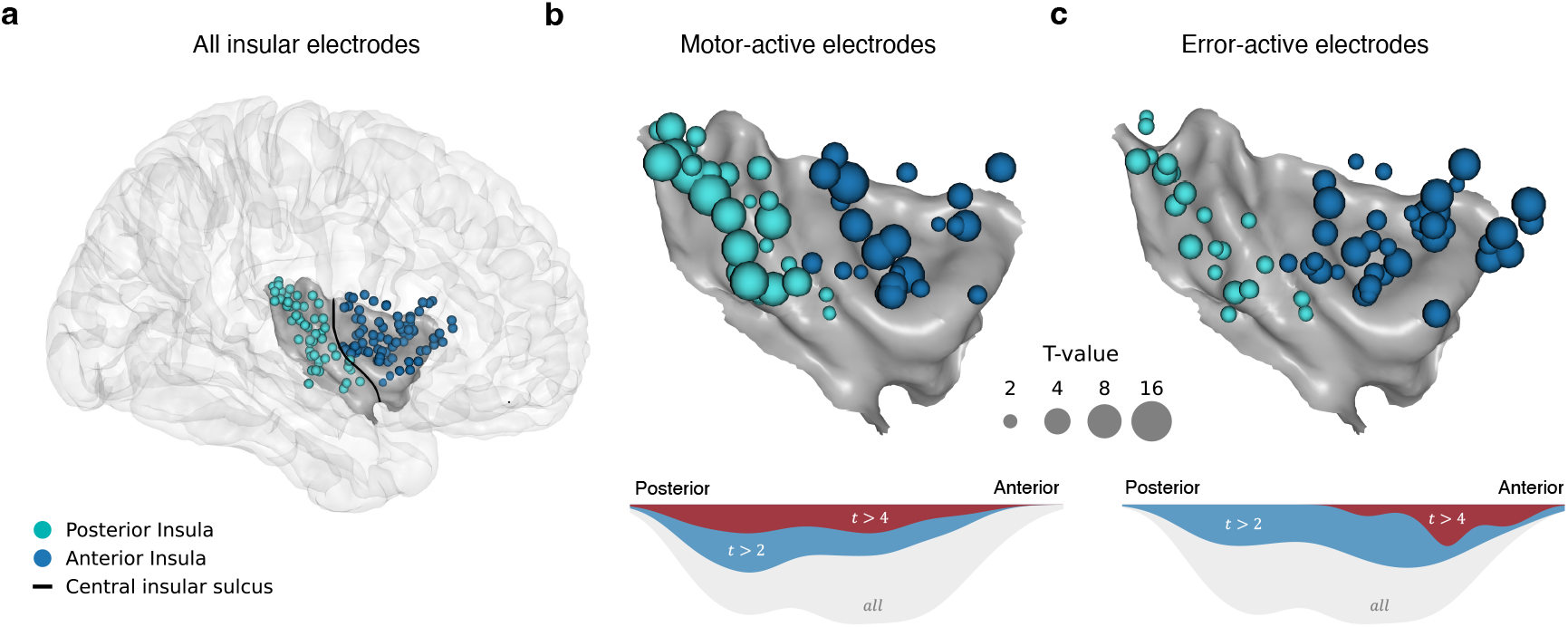
Dual role of the insula in movement control and error monitoring. **a**, Spatial distribution of all insular sites (*n* = 129) from 10 patients, warped to the MNI average brain with electrodes mirrored to one hemisphere. Electrodes were subdivided into posterior (*n* = 56) and anterior (*n* = 73) regions based on their location relative to the central insular sulcus. **b**, Movement-related insular activity during grasping (calibration phase). Responsive electrodes cluster predominantly in the posterior and central insula. The lower panel shows the smoothed distribution of movement-related electrodes along the posterior-anterior axis: all electrodes (gray), significant electrodes (*t* > 2, blue) and highly significant electrodes (*t* > 4, red). **c**, Error-related insular activity during closed-loop BCI control. Significant electrodes are more heterogeneously distributed, but highly active electrodes (red) form a distinct peak in the dorsal anterior insula.

Error-responsive electrodes during closed-loop BCI control were distributed more heterogeneously across the insula (blue distribution, Fig. 4c), with the spatial distribution revealing a higher concentration in the anterior portion. Notably, highly active electrodes (shown in red) were exclusively present in the dorsal anterior insula, forming a clear spatial peak in this region. We therefore observe that the insula serves as a heterogeneous hub, with action- and performance-monitoring signals distributed across partially different subregions.

### Real-time error decoding enables self-correcting BCI

We next asked whether error-related potentials could be transformed into actionable control signals for a self-correcting intracranial BCI. To address this, we first evaluated error decoding performance across the three target regions using a centered 500-ms sliding window applied to the subset of highly responsive electrodes (Fig. 5a). Decoding accuracy in the anterior insula and prefrontal cortex increased once the sliding window encompassed activity following a lane change. In the anterior insula, performance rose 100 ms earlier than in the PFC, consistent with its temporal lead in error processing. Peak accuracy was slightly higher in the anterior insula, reaching 91% after 625 ms compared to 90% after 710 ms in the PFC. Significance was reached 120 ms before the lane switch in the anterior insula, as the centered 500-ms window captured neural activity up to 130 ms after the switch. In the PFC, significance followed later at 180 ms post stimuli.

**Figure 5.**
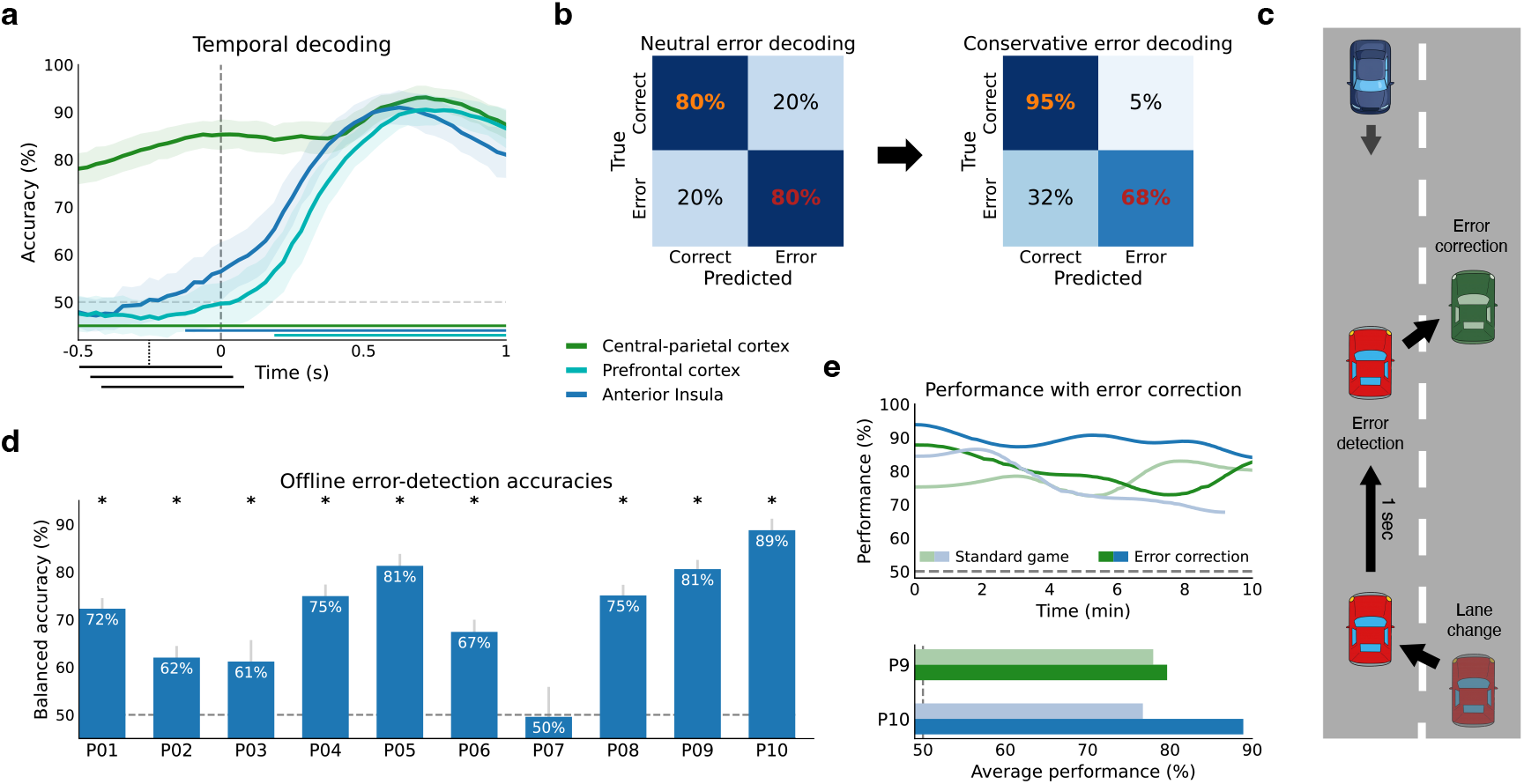
Decoding error-related activity for real-time game correction. **a**, Decoding accuracy over time across the three target regions using a centered 500-ms sliding window (mean ± s.d.). Significance is indicated by the colored bar at the bottom, aligned to object presentation (correct vs. incorrect lane switch; dark gray dashed line), with the light gray dashed line marking chance level (50%). **b**, Confusion matrices from an exemplary participant (P10) comparing neutral and conservative error decoding. In the conservative approach, a prior was applied to constrain the false-positive rate to 5% or lower. **c**, Exemplary game scenario demonstrating live error correction: the car makes an incorrect lane change, followed by error detection and subsequent correction. **d**, Offline error decoding accuracy across 10 participants, with error bars representing the standard deviation across random initializations and asterisks indicating performance significantly above chance level (50%, gray dashed line). **e**, Online performance during error correction in the last two participants. The top panel shows performance over time for the standard game and with error correction, and the bottom panel shows the corresponding average performance across the session (chance level indicated at 50%).

Temporal decoding in the central–parietal cortex displayed a distinctively different pattern, with accuracy remaining significantly above chance level (50%) throughout the entire period. Peak accuracy was reached simultaneously with the PFC at 720 ms after the stimulus, reaching 93%. This sustained performance is again consistent with the task-structure, in which central–parietal activity reflects both pre-stimulus movement-related processes and post-stimulus error-related signals.

We proceeded with decoding on the level of individual participants, where we extracted high-frequency features from the first second after lane switch to train an LDA classifier. In an example participant (P09), the model detected about 80% of errors but also misclassified 20% of correct trials, a false-positive rate too high for practical use (left confusion matrix in Fig. 5b). To improve usability, we applied a prior that shifted predictions toward correct trials, capping the false-positive rate at 5%. This adjustment preserved detection of more than 65% of errors without compromising usability for real-time BCI control (right confusion matrix in Fig. 5b).

Offline decoding across all participants exceeded chance level in 9 out of 10 participants (Fig. 5d), with performance measured using balanced accuracy to account for the unequal number of correct and erroneous trials. Decoding results ranged from 50% in the lowest-performing participant (P7) to 89% in the highest-performing participant (P10), in line with varying electrode coverage and access to error-responsive sites across participants.

Moving towards online error correction, we applied the LDA-based error model from the first closed-loop session as a second classifier running in parallel with the RNN-based movement decoder (Fig. 1a). After each lane switch, features from the first second of activity were extracted and classified to determine whether the change was erroneous or correct (Fig. 5c). When neural activity indicated an error, the car automatically returned to the original lane and blinked green for one second, providing visual feedback of the correction. The hybrid movement–error decoder was deployed in the final two participants, showing that game performance can be improved by online error correction (Fig. 5e).

In this proof-of-concept implementation, cognitive error signals were integrated into real-time device control, allowing the system to monitor its performance and autonomously correct mistakes. This demonstrates the feasibility of a self-correcting human intracranial BCI based on distributed error-related activity.

## Discussion

In this study, we leveraged intracranial electrophysiology from ten drug-resistant epilepsy patients to investigate error processing during real-time BCI use. We observed a sequential pattern of high-frequency activity following an error, beginning in the anterior insula and propagating into a broader cortical network. By sampling these signals in real time, we showed that errors can be detected and corrected with minimal delay, reducing the need for manual user intervention. Our results therefore highlight the insula as a key target for BCIs, as its dual role in movement and error processing enables the integration of action and evaluation, supporting a self-correcting human intracranial BCI.

Our calibration paradigm confirmed characteristic electrophysiological dynamics of motor activity, showing robust high-frequency increases in somatosensory cortices and corresponding low-frequency decreases spreading across adjacent regions^36–42^. We leveraged both frequency bands to rapidly train a movement decoder, enabling reliable real-time BCI control, consistent with previous studies^43,44^. Our top accuracy of about 85% likely reflects labeling limitations, as movement instructions were cued through visual prompts rather than precisely timed labels using electromyography (EMG).

Motor-related activity followed a clear temporal cascade at movement onset, beginning in the precentral gyrus and progressing to delayed responses in the postcentral gyrus and posterior insula. This sequence could reflect the hierarchical organization of motor processing, with early precentral activity supporting movement initiation, followed by sensory feedback in the postcentral gyrus, and subsequent monitoring and higher-level integration in the insula. While somatosensory activation aligns with prior intracranial studies of sequential recruitment from supplementary motor areas to primary motor and somatosensory cortices^45–49^, the role of the insula has been far less explored.

The insula’s precise contribution to movement thus remains unclear, as its concealed location within the Sylvian fissure makes it difficult to access with electrophysiological probes. Anatomical studies have demonstrated strong connectivity between the posterior insula and sensorimotor cortices, and fMRI work has linked its activity to the integration of somatosensory and interoceptive information^50–53^. However, these approaches lack the temporal resolution to capture the rapid dynamics of movement. Only a few invasive studies have addressed this question directly, reporting delayed posterior insula responses relative to primary motor cortex, with estimated latencies of 75 ms or higher^54,55^.

Extending this limited evidence, our findings highlight the delayed involvement of the posterior insula, consistent with a role in processing sensory feedback and refining motor output rather than initiating movement. We observed a delay of approximately 140 ms relative to precentral gyrus, followed by sustained activity that persisted throughout ongoing movement. We further demonstrate real-time use of insula activity for motor BCI control, underscoring its potential as a target for continuous movement decoding.

Next, we examined neural responses when the BCI failed to follow the user’s intended action. These visually perceived mismatches engaged the brain’s error network, with changes observed across the insula, prefrontal cortex and central-parietal regions. High-gamma activity first emerged in the dorsal anterior insula, followed by later engagement of the dorsolateral PFC and more widespread activity in central and parietal cortices. This pattern might reflect a temporal progression from error detection in the insula, to higher-order cognitive evaluation in the PFC, and ultimately to updating sensorimotor plans in central–parietal regions.

Our observation of early activity in the anterior insula aligns with its central role within the salience network, where it is thought to detect behaviorally relevant events and signal the need for adaptive control^11,13,16,17^. Beyond detection, the anterior insula may coordinate with other brain regions to initiate compensatory actions and maintain goal-directed behavior. Evidence points to feedforward interactions with regions including the PFC, premotor cortex and anterior cingulate cortex, observed from selected pairs of electrodes^14,16,50,51^. Recent work using neuroimaging and focused ultrasound stimulation further maps a pathway of error processing from visual areas to the anterior insula, which then continues to prefrontal and motor regions^56^. Our findings extend this framework by demonstrating consistent population-level delays across these regions, identifying their precise spectral contributions in the gamma/high-gamma range, and translating these signals into real-time error correction during BCI control.

While error-related activity was most strongly observed in the anterior insula and prefrontal cortex, central regions likewise exhibited neural responses. These signals remain more challenging to interpret due to their strong involvement in movement, functionally overlapping with error processing. In our task, central activity differentiated correct from erroneous trials even before the error occurred, reflecting a task limitation where correct trials involved pre-movement while erroneous trials did not. Still, we find error-related signals following the stimulus, consistent with prior studies of human motor cortex^28,29^, although care must be taken to ensure that decoders do not simply capture compensatory movements. Disentangling these overlapping processes is particularly challenging in naturalistic environments but may be achieved using characteristic activation profiles and their lower-dimensional embeddings.

After characterizing the error network, we subsequently sampled these signals in real time to enable error correction during ongoing BCI control. By running an error decoder in parallel with the movement decoder, the system was able to detect and reverse unintended actions, demonstrating a human intracranial BCI that can correct its own mistakes. Importantly, we relied on error signals from the insula and prefrontal cortex, key regions for cognitive control, rather than from the motor cortex, where activity may be confounded by compensatory movements. This approach highlights the potential of higher-order cognitive regions as novel BCI targets, paving the way for interfaces that can monitor, evaluate, and correct their own actions.

Previous work on error processing during BCI control has mainly focused on offline classification of error-related potentials, mostly using motor cortical signals in humans and non-human primates^27–29^. Active, real-time error correction has so far been limited to non-invasive EEG studies^30–32^ and non-human primates, where incorporating error signals improved bit rate and reduced error rates during cursor control^33^. Here, we extend this body of work by demonstrating online error correction in humans using intracranial recordings and sampling directly from higher-order cognitive regions.

Although this study contributes to the development of cognitively enhanced BCIs, several limitations remain. First, the error decoder operated only on end-of-trial signals, preventing continuous tracking of errors with varying magnitude. Second, electrode coverage did not sufficiently sample the anterior cingulate cortex, a region strongly implicated in error processing and performance monitoring^9,10,57^, resulting in a less complete view of the error network. Third, online error correction was implemented late in the study and therefore tested in only two participants, making it a proof-of-concept rather than a full cohort evaluation. Fourth, experiments were carried out in a controlled laboratory environment, leaving translation to real-world BCI use addressing error timing and strength an open challenge. Future work should focus on developing continuous error decoders that capture error magnitude, while sampling distributed nodes of the network in everyday settings.

Our findings bear important implications for the development of BCIs that go beyond decoding motor intentions. By accessing higher-order brain regions such as the prefrontal cortex and insula, these systems could detect internal states related to error awareness, uncertainty, or conflict. Such an interface would infer a user’s abstract cognitive processes, potentially enabling more naturalistic and adaptive interactions aligned with the individual’s goals. This work opens the door to next-generation BCIs that directly engage with a user’s cognitive state, while also raising critical questions about autonomy, privacy, and the broader societal impact of these technologies.

## Methods

### Participants

Data was recorded from ten patients with drug-resistant epilepsy (6 female, age range 26-56 years) undergoing clinical monitoring to localize their epileptic seizure zone. Patients underwent implantation of stereotactic electroen-cephalography (sEEG) electrodes, with electrode placement solely based on clinical needs. Surgical implantation took place in the Maastricht University Medical Center (MUMC), the Netherlands, with subsequent monitoring for about two weeks in the Academic Center for Epileptology, Kempenhaeghe. Before the experiment, each patient provided their written informed consent to voluntarily participate in research as well as subsequent data usage. Experimental paradigms were in accordance with protocols of the institutional review board of Maastricht University and Epilepsy Center Kempenhaeghe (METC 20180451).

### Intracranial recordings

Neural recordings were obtained using Microdeep intracerebral electrodes (Dixi Medical, Beçanson, France) containing 5-18 recording channels per shaft (2 mm long contacts, 1.5 mm intercontact distance, 0.8 mm shaft diameter). Data was recorded using two to three Micromed SD LTM amplifiers (Micromed S.p.A., Treviso, Italy), each sampling 64 channels with a sampling rate of 1024 Hz. Each participant’s electrodes were referenced to a common white-matter contact showing no epileptic activity. Neural data and behavioral markers were synchronized using Lab Streaming Layer (LSL)^58^.

### Anatomical electrode localization

A presurgical T1-weighted magnetic resonance imaging (MRI, 3 Tesla) scan and a postsurgical computed tomography (CT) scan were obtained for each patient to identify electrode locations within the brain. Freesurfer (https://surfer.nmr.mgh.harvard.edu/) was used to extract the pial surface from the MRI, enabling co-registration of electrode locations with each patient’s brain model. Cortical anatomical labels were derived using the Destrieux atlas^59^ through a custom version of the semi-automated Python package *img_pipe*^60^. Electrode locations were then warped to the MNI152 average brain for visualization. Finally, electrode labels for the insula were manually refined in each participant’s personalized brain model in consultation with a neurosurgeon, distinguishing posterior and anterior regions (Fig. 4a). In addition to the insular sites, one adjacent white matter contact was included to capture activity from a broader region. This resulted in a total of 1,317 electrodes across the whole brain (Fig. 1b), 129 of which were located in and immediately around the insula.

### Experimental paradigm

Patients performed the task while seated comfortably in a chair in their private hospital room, with all instructions explained in Dutch by one of the researchers. The experiment began with a calibration session, during which patients repeatedly grasped with both hands following move/rest cues displayed on a screen (Fig. 1c). Each trial lasted 2.5–3.5 seconds, with the exact duration randomized to prevent anticipation, and a time bar providing visual feedback for more precise timing. Using the supervised data from this session, we trained a decoder that continuously estimated movement probability. Subsequently, patients completed a brief two-minute neurofeedback session, viewing a real-time graph showing the decoder’s predictions over the last 10 seconds to familiarize themselves with how the system interpreted their movements.

Next, patients engaged in a closed-loop car game lasting 5–10 minutes to evaluate decoder performance and collect data for error potential training (Fig. 1e). The game was developed in Pygame and featured the control of a red car on the screen, which switched lanes whenever the decoder’s movement probability exceeded a predefined threshold. The blue car, acting as an obstacle, appeared from the top of the screen, giving patients around 3-4 seconds to react and avoid a collision by switching lanes. After each lane change, the red player car was locked for 3 seconds to prevent rapid consecutive actions. Data from this session was used to train an error decoder to distinguish between correct lane changes (avoiding the obstacle) and erroneous ones (moving into the obstacle lane). In a subsequent session, this error decoder was deployed for real-time error correction during gameplay.

Game performance was quantified as the percentage of obstacles successfully passed. Obstacles appeared in one of two possible positions and were presented in a pseudo-randomized order, keeping the number of occurrences balanced between positions, resulting in a chance level of 50%. Performance over time was calculated using a rolling average over 20 trials, with each point representing the percentage of correctly passed obstacles within that window (Fig 2g). The resulting trace was further smoothed using a Gaussian kernel.

### Data preprocessing

Neural data was preprocessed for offline analysis using a custom Python pipeline designed to reduce noise and estimate local activity. First, the raw iEEG recordings were re-referenced using an Electrode Shaft Reference (ESR)^61^. In this step, the average activity across all electrodes on a given shaft was subtracted from each electrode at every time point, thereby reducing shaft-wide artifacts and isolating local neural signals. Next, the signals from each electrode were normalized (z-scored) across the entire experimental session, resulting in a cleaned version of the raw data.

To further prepare the data for analysis, we applied a series of filtering and power estimation steps. First, the data was band-pass filtered into frequency ranges of interest, isolating high-frequency components (60–200 Hz) and low-frequency components (8–30 Hz) relevant for motor decoding. Importantly, we used zero-phase, non-causal Butterworth filters applied in a two-pass process, ensuring that no phase shifts were introduced by the filtering. Next, the power of each band was obtained by calculating the signal envelope using the absolute value of the Hilbert transform. Finally, the resulting power signals were smoothed using a centered 250 ms Gaussian kernel and normalized (z-scored) once more to produce standardized power values for each channel.

### Real-time data handling

During closed-loop BCI control, neural data arrived in small, irregular chunks as it was streamed from the acquisition system. Each chunk was immediately re-referenced and normalized using the mean and standard deviation obtained from the calibration session. The processed chunk was then appended to a 4-second circular buffer, which was continuously updated to always contain only the most recent data.

Every 125 milliseconds, the full buffer was preprocessed to extract features for real-time decoding. The buffer was first filtered with zero-phase filters to isolate relevant frequency bands and prevent edge artifacts. Next, the instantaneous power of each band was then computed by taking the absolute value of the Hilbert transform, followed by channel-wise normalization using the statistics from the calibration session to ensure consistency with the training data. Lastly, the latest 250 ms segment was extracted as the prediction window, which was averaged across time to obtain one feature per channel and band.

### Movement decoder design, training and usage

The movement decoder was trained on approximately three minutes of labeled data from the calibration session. Data was segmented into non-overlapping 250 ms windows, with each window labeled by the majority class within it and kept in temporal order.

For each patient, the feature set consisted of two features per channel, capturing both high-frequency activity and low-frequency activity. We applied feature selection by calculating the F-score between each feature and the calibration labels, which quantifies how well a feature separates the two classes. Features with an F-score above 10 were retained, with a minimum of 20 and a maximum of 100 features used for training per participant.

We used a recurrent neural network as movement decoder, designed to capture temporal dependencies in the neural features. The network consisted of an input layer corresponding to the selected features, followed by two LSTM layers with 10 units each, and a final output layer with a single unit estimating the continuous movement probability. In addition, a learnable skip connection was included between the input and output layers, which was weighted together with the LSTM output to form the final prediction. This design allows the decoder to merge immediate neural responses with longer-term temporal dependencies. Lastly, we set a prior towards resting to reflect the higher prevalence of rest during closed-loop control, ensuring that the decoder remained robust to this natural imbalance.

The decoder was trained for 60 epochs with a batch size of 32 using the Adam optimizer. The LSTM layers and the skip connection were optimized jointly, with the skip connection using ReLU units. To reduce overfitting, we applied dropout and gradually decreased the learning rate during training using a scheduler.

Performance was evaluated using 10-fold cross-validation, keeping the temporal order of windows intact. All steps, including feature selection, were performed only on the training data within each fold to prevent data leakage. After cross-validation, a final decoder was trained on the full dataset using the same procedure and saved for closed-loop control.

To assess how the amount of training data affects decoder performance, we repeated the analysis for each participant while gradually increasing the size of the training set (Fig. 2e). Training began with an untrained decoder (0 samples) and increased in 5-second increments during the first minute and 20-second increments thereafter. Twenty percent of the data were reserved for testing, and the remaining data was balanced between classes and kept in temporal order to preserve the sequential structure of the recordings. This procedure was repeated 50 times with different starting positions and performance was averaged across repetitions to reduce variability.

During real-time BCI operation, preprocessed features were sent to the decoder every 125 milliseconds. The decoder applied the previously selected feature set and generated movement probability predictions, with each prediction updating the RNN’s internal state. A movement command was issued only when the predicted probability of movement exceeded 70% for at least three consecutive windows, ensuring stable detection and reducing false positives. Once triggered, the command was converted into game actions and transmitted via Lab Streaming Layer to the game, which ran in parallel with the decoder and the feature extraction pipeline.

### Analysis of movement-related activity

To identify movement-related activity, we segmented the calibration data into non-overlapping 250-ms windows and computed the average spectral power for each segment. Labels were assigned based on whether most time points within the segment occurred during movement or rest. For each electrode, we compared the average power between movement and rest using a two-sample Welch’s t-test, which accounts for potential differences in sample size and variance. The resulting t-values were used as a measure of movement-related modulation, whereas electrodes with (*t* > 4) were considered motor-relevant, reflecting strong and reliable activity rather than subtle differences (Fig. 2c).

To examine the temporal dynamics of movement initiation, we analyzed high-frequency activity across three regions of interest: the precentral sulcus (including primary motor cortex, M1), postcentral sulcus (including primary somatosensory cortex, S1), and posterior insula. Neural signals were aligned to the prompted movement onset, first averaged across electrodes per region and patient, and then averaged across trials to generate time-resolved activity profiles (Fig. 2d). These profiles were baseline-corrected using the interval from *−*2 to *−*0.5 seconds prior to movement onset, with the average activity during this resting period defined as zero. Deviations from baseline were then assessed at each time point with a one-sample t-test.

### Analysis of error-related activity

Next, we examined error-related activity by comparing correct lane changes, where the car moved away from an incoming obstacle, with erroneous lane changes, where it moved into the obstacle lane (Fig. 3a). High-frequency activity was analyzed during the first second after the lane change in three target regions: the anterior insula, prefrontal cortex, and central-parietal cortex. The central region included the pre- and postcentral areas as well as the supplementary motor area, which was grouped with the parietal region due to similar patterns of activity observed during the task.

We quantified differences in activity between conditions by calculating t-values for each electrode. To account for variability in response timing, three 500-ms windows were defined (0–500 ms, 250–750 ms, and 500–1000 ms) and examined using a two-sample t-test. The window with the highest absolute t-value was selected for each electrode to represent the strongest modulation, independent of its precise timing. Consistent with earlier analysis, highly responsive electrodes were defined as those with t-values greater than 4, marking channels with clear and robust error-related activity.

Subsequently, we obtained temporal activity profiles for each region of interest by examining responsive electrodes over time. To create a local estimate of population activity, we first averaged HFA across electrodes within each region and participant. This step preserved individual trials while reducing variability introduced by differences in electrode counts, preventing participants with many electrodes from disproportionately influencing the group signal. As a result, the regional signal for each participant reflects coordinated activity across electrodes rather than being dominated by isolated, highly active sites. Finally, these trial-level signals were stacked across participants and averaged to generate group-level activity profiles (Fig. 3c).

Next, we compared high-frequency activity during correct and erroneous trials across all electrodes in each region of interest (Fig. 3d). Because electrodes contained a different number of trials, and each participant contributed a varying number per region, we used a linear mixed-effects model to account for these differences. The model included random intercepts for electrodes nested within participants, allowing condition effects to be estimated while controlling for inter-individual and electrode-specific variability (implemented in Python using statsmodels, with trial condition as the dependent variable and participants and electrodes as random factors).

To better capture temporal variability at the electrode level, we assessed the peak latency for each highly responsive electrode during erroneous trials. HFA was averaged across these trials to generate a representative activity profile for each electrode, which typically exhibited a distinct peak compared to weakly responsive sites (Fig. 3e). Peak latency was then defined as the time point corresponding to the maximum response within this profile (Fig. 3f). Differences in peak latency between regions were tested using pairwise Mann–Whitney U tests, which do not assume a specific underlying data distribution and can accommodate unequal numbers of electrodes across regions. Finally, we characterized the precise spectral profile of the error response at both the single-electrode and group level. For the single-electrode analysis, we computed a spectrogram for each correct and erroneous trial, which was subsequently smoothed in both the time and frequency domains using a Gaussian kernel. Differences between erroneous and correct trials were then quantified by calculating a t-value for each time–frequency bin, with significant clusters of diverging activity (*p* < 0.05) being outlined (Fig. 3h).

For the group-level analysis, we first established a baseline frequency distribution by computing the power spectrum across the entire experimental session for each electrode. Power spectra were then calculated for the one-second window following an erroneous lane change, which were inherently noisy due to the short segment length. To obtain stable estimates, these spectra were smoothed using a Gaussian kernel and averaged across trials for each electrode. Subsequently, each frequency bin was expressed as the percentage change in power relative to the baseline distribution obtained earlier. The resulting normalized spectra were averaged across all highly responsive electrodes in all target regions, producing the final group-level frequency profile of error-related activity (Fig. 3g).

### Decoding error responses

To investigate the predictability of error responses over time, we examined decoding performance of HFA for each target region. First, all electrodes were aligned to the time of the lane change and stacked within their respective region to form a region-specific dataset. To account for the different number of trials across electrodes, we selected the minimum number of trials present in any electrode, with an equal split between correct and erroneous trials for each channel. This process was repeated 100 times with different subsets of trials to include all data and account for variability in trial selection. The results were subsequently averaged across repetitions, enabling trial-based decoding using all electrodes in a region as features (Fig. 5a).

Having prepared the dataset, we next partitioned the neural activity into 500 ms centered windows, shifting them by 30 ms over a 2-second period. Within each window, high-frequency activity was averaged across time for every electrode to generate features for classification. Prediction was performed using a multilayer perceptron (MLP) with a single hidden layer of 10 units and L2 regularization to promote feature diversity. This approach encouraged the model to rely on contributions from all electrodes rather than overfitting to a few highly informative ones. Stratified 10-fold cross-validation was used to maintain balanced classes in each fold, while decoding performance was assessed as the average classification accuracy across folds. Lastly, we tested for significance using a Monte Carlo approach by shuffling trial labels and repeating the decoding 100 times, with the 95th percentile of the resulting accuracy distribution serving as the significance threshold.

Moving from group-level decoding to participant-specific analysis, we trained error decoders for each individual to distinguish correct from erroneous trials using data from a single closed-loop session. Using all electrodes from a participant as features, we trained a linear discriminant analysis (LDA) classifier on data from the first second after each lane change. LDA was chosen because it can be rapidly trained and modified, allowing for quick adjustments and setting of a prior. The classifier’s decision threshold was iteratively adjusted by shifting the prior toward correct lane changes until the false positive rate was reduced to 5% or lower (Fig. 5b). This process was repeated 10 times for each participant to account for variability in initialization, with decoding performance quantified using balanced accuracy to address the unequal number of correct and erroneous trials (Fig. 5d). Lastly, significance was determined using a Monte Carlo approach as described earlier.

### Literature research

Literature research was conducted using Google Scholar, PubMed, and ResearchGate, resulting in the inclusion of 61 relevant papers. Of these, 12% had a female first author and 10% had a female last author.

## Acknowledgements

P.W., M.V., P.K. and C.H. thank the INTENSE consortium for their funding (project number 17619 of the research programme NWO Crossover Programme, which is partially financed by the Dutch Research Council NWO).

## Author contributions statement

P.W., A.C., J.P.v.D. and M.V. recorded the data. P.W., C.H., M.C.O., Y.T., P.L.K. designed the experiments. P.W., S.G., L.O., C.H. analyzed the data. P.W. wrote the first draft of the manuscript. All authors contributed to the final version of the manuscript.

## Additional information

### Data and code availability

In line with an open science approach, we are publicly sharing all data (https://osf.io/wqsgr) and code (https://github.com/paulweger/InsularErrorNetwork.git) to recreate the results.

### Consent to publication

All authors have read and approved the final manuscript and consent to its publication.

